# Can music modulate gene expression involved in traumatic brain injury? An integrative transcriptomic and epigenomic proof of concept

**DOI:** 10.1101/2025.07.02.662815

**Authors:** Alberto Gómez-Carballa, Sara Pischedda, Nour El Zahraa Mallah, Wiktor Nowak, Sensogenomics Working Group, Federico Martinón-Torres, Laura Navarro, Antonio Salas

**Affiliations:** Genetics, Vaccines and Infections Research Group (GenViP), Instituto de Investigación Sanitaria de Santiago, 15706 Universidade de Santiago de Compostela, Santiago de Compostela, Galicia, Spain; Unidade de Xenética, Instituto de Ciencias Forenses, Facultade de Medicina, Universidade de Santiago de Compostela, and Genética de Poblaciones en Biomedicina (GenPoB) Research Group, Instituto de Investigación Sanitaria (IDIS), 15706 Hospital Clínico Universitario de Santiago (SERGAS), Galicia, Spain; Centro de Investigación Biomédica en Red de Enfermedades Respiratorias (CIBER-ES), Madrid, Spain; Translational Pediatrics and Infectious Diseases, Department of Pediatrics, 15706 Hospital Clínico Universitario de Santiago de Compostela, Santiago de Compostela, Galicia, Spain

**Keywords:** Brain Damage, Acquired Brain Injury, Traumatic Brain Injury, Transcriptomics, Gene Expression, RNA-seq, Epigenomics, microRNA

## Abstract

Music-based interventions have shown promise in aiding recovery in individuals with brain damage (BD), including acquired and traumatic brain injury (ABI and TBI). To investigate the underlying molecular mechanisms, we conducted a proof-of-concept study integrating transcriptomic and epigenomic data. We compared gene expression changes driven by musical stimuli in healthy individuals exposed to music with transcriptomic data from TBI patients, identifying 154 common differentially expressed genes (DEGs) affected in TBI and impacted by music, including *STK35*, *HNRNPA0*, *MT-CYB*, and *MT-ND2*. These genes are notably involved in mitochondrial function, synaptic plasticity, and neurodegenerative processes. Epigenomic analysis revealed that 92 of these DEGs also exhibited significant differential methylation in TBI samples, particularly in genes associated with neurodevelopment, neuroinflammation, and mood regulation. Pathway enrichment and cross-validation in independent datasets of TBI patients highlighted the cGMP– PKG signaling pathway, a key regulator of memory and plasticity. Co-expression network analysis further revealed six gene modules significantly associated with TBI, including modules centered on *TXNIP* and *ALG13*, genes previously implicated in neuroinflammation and epilepsy, two conditions commonly associated with TBI. Interestingly, several hub genes from these modules also showed sensitivity to musical stimuli, suggesting that music may target biologically relevant networks involved in TBI pathology. These findings provide preliminary evidence that music exposure may modulate gene expression and methylation in ways that support neural repair and functional recovery. This molecular insight reinforces the potential of music as a biologically grounded component of neurorehabilitation. Future studies should directly assess transcriptomic and epigenomic responses to music-based interventions in BD patients.

## Introduction

Music is one of the most universal and complex forms of human expression, deeply embedded in all cultures and stages of life [1]. While its psychological and social benefits have long been recognized, the biological mechanisms underlying the effects of music on the human brain are only beginning to be understood. Recent developments in genomics and systems biology offer powerful tools to uncover these mechanisms, particularly through transcriptomic and epigenomic analyses.

In the past two decades, cognitive and neuroscientific research has underscored the role of music in shaping brain function and structure, highlighting its capacity to engage extensive neural networks beyond the auditory cortex [2-5]. Parallel to these neurophysiological insights, genetic studies have begun to identify loci and heritable traits associated with musical ability and responsiveness [6, 7]. A particularly promising development is the emergence of sensogenomics, a field that explores how sensory stimuli such as music influence gene expression and other molecular processes [8]. Within this framework, transcriptomic approaches, especially RNA sequencing (RNA-seq), have proven instrumental in revealing the gene networks modulated by musical exposure not only in healthy individuals but also in patients affected by different cognitive conditions [8-10].

The sensogenomics project (www.sensogenomics.com/en/; [8-10]) has pioneered the use of transcriptomic analysis to explore how music affects gene expression in various physiological and pathological contexts. In a landmark study, researchers demonstrated that listening to music altered the transcriptome of healthy individuals and identified genes implicated in synaptic plasticity, mitochondrial activity, and neuroprotection. Further, these transcriptional responses were found to intersect with pathways relevant to neurodegenerative conditions such as Alzheimer’s disease, illustrating the potential of music to modulate disease-relevant molecular circuits [8-10].

However, omics-based research remains scarce in the context of brain damage (BD) caused by stroke, traumatic brain injury (TBI), or other acquired brain injuries (ABI), despite mounting evidence supporting the clinical efficacy of music in neurorehabilitation [11]. While previous studies have explored behavioral and functional outcomes of music-based interventions in BD [12-17], few have examined how these interventions might influence gene regulatory mechanisms. Notably, recent efforts have started to integrate epigenomic data, such as DNA methylation patterns, with transcriptomic findings to understand how music might exert longer-term regulatory effects on the genome, especially in relation to neural repair and inflammation [9, 10].

Given the urgent need for novel, non-pharmacological interventions in the treatment of BD, the application of transcriptomic and epigenomic tools could yield critical insights into the biological basis of music’s therapeutic potential. Such insights could help identify biomarkers of response to music-based therapy and guide the development of precision interventions tailored to individual molecular profiles.

In this context, the present study aims to advance the field by investigating how music exposure influences gene expression and DNA methylation in BD. We conducted a proof-of-concept integrative analysis, combining transcriptomic and methylation data from TBI patients with expression profiles from healthy individuals exposed to music [9]. Our approach provides a first step toward understanding the molecular pathways that may mediate the beneficial effects of music in neurorehabilitation, setting the stage for future studies directly targeting transcriptomic and epigenomic changes in TBI patients undergoing music-based interventions.

## Methods

### Transcriptomic Analysis

Gene expression data related to TBI were retrieved from the Gene Expression Omnibus (GEO) database, including two datasets: *i*) GSE254880, which includes RNA-seq data generated from serum-derived neuronal exosomes of patients with severe TBI (*n* = 8) and healthy controls (*n* = 8); and *ii*) GSE131695, comprising serum miRNA profiles from acute TBI patients collected at 24, 48, and 96□hours post-injury (*n* = 33), along with age-matched healthy controls (*n* = 6), as well as chronic TBI patients (*n* = 6) and their respective controls (*n* = 6).

Raw RNA-seq data from GSE254880 were downloaded from the Sequence Read Archive (SRA) database and processed. One healthy control sample was excluded because only one of the two files from the paired-end sequencing data was provided, resulting in *n* = 8 TBI and *n* = 7 healthy controls. Quality control of FASTQ files was conducted using *FastQC* [18]. Adapter removal, trimming, and QC filtering of reads were performed using *trimmomatic* [19]. Filtered sequences were mapped against the reference human genome (GRCh38) using the aligner *STAR* [20]. Counts generated from *STAR* were used for downstream analysis. Raw count data from the miRNA GSE131695 dataset were directly downloaded from GEO. Only the 24-hour post-injury data were used from the acute TBI samples (*n* = 21) for the analysis. Lowly expressed genes (dataset GSE254880) and miRNAs (dataset GSE131695) were filtered out through the *filterByExpr* function from the *EdgeR* package [21]. Data normalization was performed with the *DESeq2* [22] and *RUVSeq* [23] packages. Reference genes and miRNAs for the *RUVg* function were detected by selecting invariantly expressed genes obtained from a naïve differential expression analysis.

To predict gene targets of the differentially expressed miRNAs in the GSE131695 dataset, we used the *miRNAtap* R package [24]. Predictions were based on five databases: *PicTar*, *DIANA*, *TargetScan*, *MiRanda*, and *miRDB*. Genes were ranked using a geometric mean score, and a gene was retained as a target if it was predicted by at least three of the five databases.

We performed an over-representation analysis (ORA) approach to identify biological processes related to the genes affected by the musical stimuli from the study by Gómez-Carballa et al. [9] (*P*-value <0.05) that were also found to be dysregulated in the GSE254880 TBI dataset. ORA was conducted using the *clusterProfiler* [25] R package and gene sets from the C5 ontology collection in the Molecular Signatures Database (*MSigDB*), which includes Gene Ontology (GO) terms and *Human Phenotype Ontology* (HPO) categories, as well as the *Kyoto Encyclopedia of Genes and Genomes* (KEGG). We applied the Benjamini-Hochberg procedure to correct for multiple testing, both *P*-value and *Q*-value thresholds were set at 0.05. The analysis was restricted to gene sets with a minimum size of 20 and a maximum of 550 genes.

The *GSVA* R package was used to calculate pathway activity scores using its internal *GSVA* method [26].

To assess statistical significance between groups, we computed a Wilcoxon rank-sum test. Heatmaps and upset plots were generated using the *ComplexHeatmap* [27] and *ComplexUpset* [28] R packages, respectively.

### DNA methylation analysis

In our search for existing datasets on brain damage/injury and epigenetics, we identified four relevant studies with publicly available DNA methylation data in the GEO. Three of these epigenome-wide association studies have examined DNA methylation patterns in ischemic stroke patients, highlighting associations with stroke outcomes [29], biological aging [30], and broader epigenetic alterations following cerebrovascular injury [31] (GEO accession numbers: GSE203399, GSE69138, and GSE280206, respectively). However, these datasets only provided methylation data from stroke patients and lacked appropriate control groups, which limited their suitability for our analysis. Therefore, we focused on a fourth dataset (GSE155426), which provided DNA methylation data from a cohort of active-duty military personnel, including individuals with and without a history of TBI [32]. We used only baseline samples collected before blast exposure (*n* = 33). Participants were categorized into two groups based on self-reported TBI history: “YES” for those with a lifetime history of TBI, and “NO” for those without. Raw IDAT files generated using the Illumina MethylationEPIC BeadChip array (from whole blood), were imported into R and processed using standard quality control and preprocessing steps. Low-quality probes were removed, and the data were normalized to obtain β-values, which represent the proportion of DNA methylation at each CpG site. Differential methylation between individuals with and without a history of TBI was assessed using independent t-tests. To explore the discriminative potential of the identified CpGs, PCA and heatmap were generated to visualize sample clustering by TBI history. In addition, boxplots were generated to display the distribution of methylation levels at the top CpG sites identified through the contrast analysis. ORA was performed using the *clusterProfiler* R package, incorporating both GO and KEGG databases.

### Co-expression Module Analysis

To investigate gene co-expression patterns potentially influenced by TBI, we applied the weighted gene co-expression network analysis (WGCNA) approach in R to the dataset GSE254880. To enhance signal specificity, we selected the top 25% most variable genes across all samples (*n* = 4,376). Considering the sample size of the dataset, a recommended soft-thresholding power of 18 was chosen to ensure approximate scale-free topology. We then computed the topological overlap matrix (TOM) and its associated dissimilarity (1 – TOM), followed by hierarchical clustering. Co-expression modules were identified using a minimum module size of 30 genes and were merged using a height cutoff of 0.25. Each module was assigned a distinct color label. To examine associations of the modules with between modules and TBI, module eigengenes were correlated with the sample trait (control *vs*. TBI comparison). Modules significantly associated with TBI were prioritized based on gene significance (GS), and key hub genes were determined using module membership (MM), reflecting their intramodular connectivity. Modules were named according to their respective hub genes. Finally, to interpret biological functions, we performed enrichment analysis using the *compareCluster* function from the *clusterProfiler* R package, focusing on Gene Ontology (GO) terms for biological processes.

## Results

### Gene expression, music, and traumatic brain injury

We aimed to infer the potential effect of musical stimuli on the transcriptome of TBI patients through a comparative strategy, as a preliminary step towards elucidating the potential of music-based therapies influencing gene expression in TBI. For this purpose, we first downloaded, processed, and analyzed the transcriptomic profiles of neuron-derived exosomes isolated from the serum of severe TBI patients by comparing them with those from healthy controls. Subsequently, we compared blood gene expression changes induced by musical stimuli in healthy controls from Gomez-Carballa et al. [9] with these transcriptomic data obtained from patients with severe TBI and controls.

TBI patients showed characteristic transcriptomic profiles different from those observed in healthy donors (**Figure 1A**). Thirty-one DEGs emerged as the most relevant genes related to the TBI condition (adjusted P-value < 0.05), clearly separating TBI and control gene expression profiles (**Table S1**, **Figure 1B**). Twenty-one genes showed upregulation (68%), whereas only ten genes were found to be downregulated in TBI cases. This pattern towards the upregulation in TBI was also observed among the top 10 ranked DEGs (**Figure 1C**). Notably, two of the top dysregulated genes were mitochondrial genes, *MT-CYB* and *MT-ND2*, highlighting the importance of mitochondrial dysfunction in TBI.

**Figure 1.**
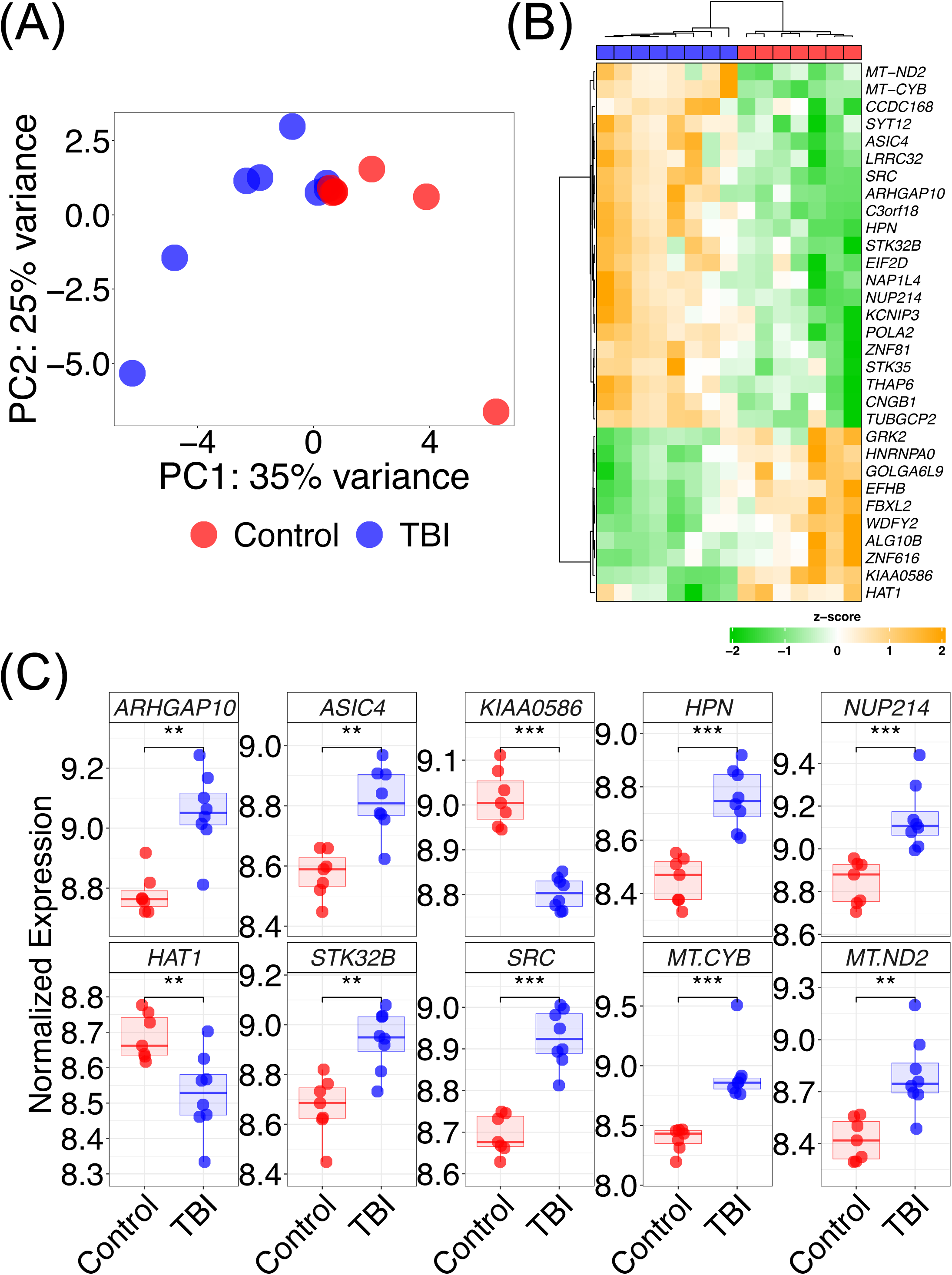
Transcriptomic analyses reveal shared and divergent expression patterns between music-altered genes and genes dysregulated in BD. (A) PCA plot of gene expression profiles of TBI cases and healthy controls from the GSE254880 dataset. (B) Heatmap representing DEGs (adjusted *P*-value < 0.05) between TBI cases and healthy controls from the GSE254880 dataset. (C) Boxplots of the top 10 DEGs (adjusted *P*-value < 0.05), ranked by Wald statistic (*stat*) column obtained from the differential expression analysis (GSE254880 dataset). Two-sided Wilcoxon *P*-values are displayed; asterisks indicate statistical significance (*** *P*-value < 0.001, ** *P*-value < 0.01).

From the previously published list of DEGs associated with musical stimuli [9], 154 were also significantly altered in TBI, with 87 showing the same direction of expression change and 67 displaying an opposite pattern (**Figure 2A**; **Table S2**). These opposite expression changes between musically stimulated donors and TBI could be indicative of a compensatory effect of musical stimuli on these TBI-related genes. *STK35* and *HNRNPA0* were among the most significantly dysregulated genes in TBI (**Figure 2A**). The expression profiles of these two gene sets can effectively distinguish severe TBI patients from controls, suggesting that these genes altered by musical stimuli may also play a role in TBI pathophysiology (**Figure 2B**).

**Figure 2.**
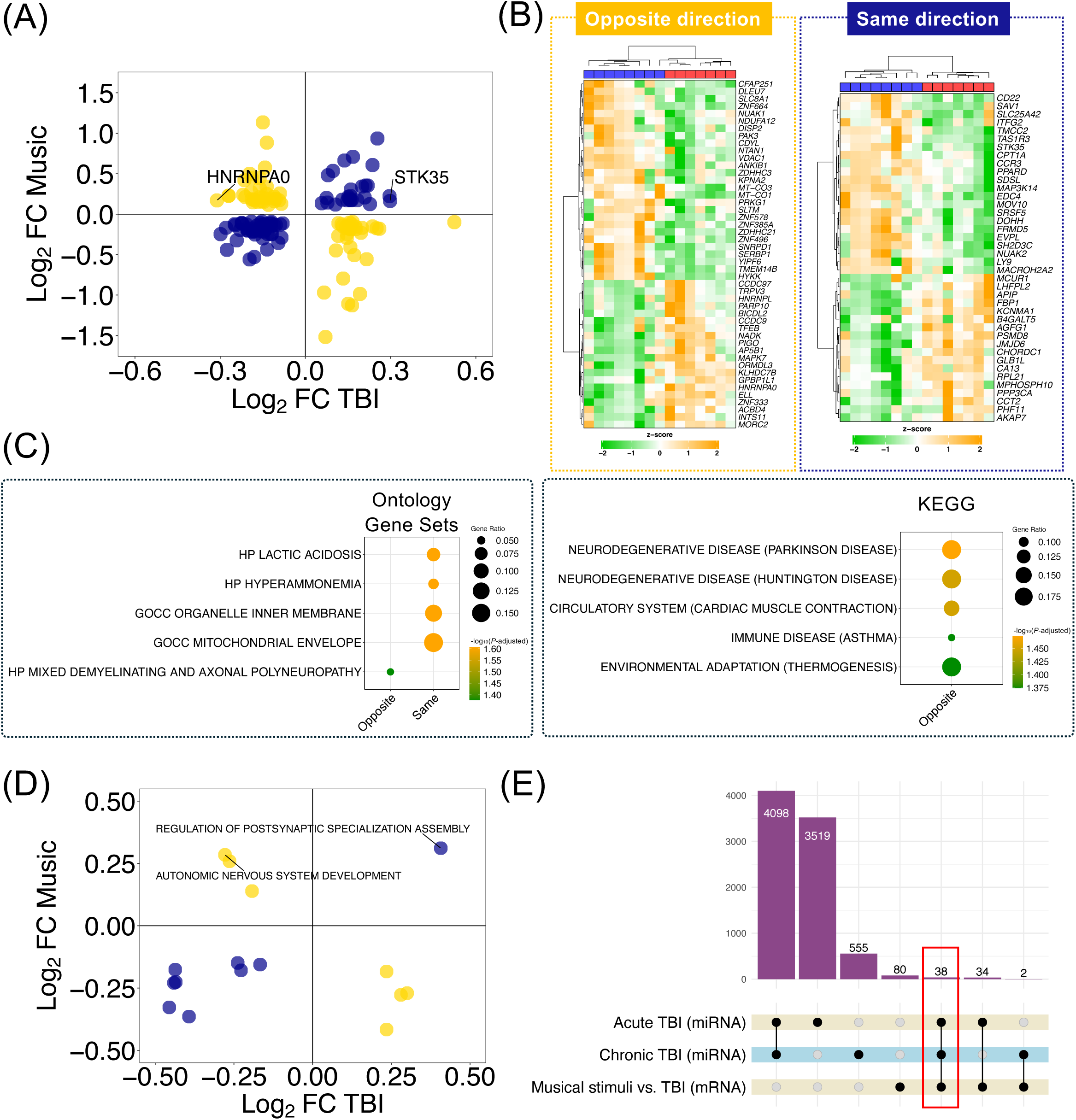
Comprehensive comparison of gene-and pathway-level expression changes between those induced by musical stimuli and those associated with brain damage. (A) Correlation plot of the log_2_FC values for 154 genes significantly altered by musical stimuli [9] and TBI. Genes with concordant expression direction are shown in dark blue; those with opposite patterns are shown in yellow. Genes significantly dysregulated in TBI (adjusted *P*-value < 0.05) are labelled. (B) Heatmap comparing the expression profiles of genes showing opposing (left) and concordant (right) expression changes in response to musical stimuli in healthy donors (compared to non-stimulated controls) and in TBI patients (compared to healthy controls). (C) Bubble plot showing the enriched terms (adjusted *P*-value <0.05) detected in the over-representation analysis from the clusters of genes showing opposite and same expression changes in TBI and musically stimulated donors. Point size indicates gene ratio (defined as the number of genes in the input list that are annotated to the corresponding term/number of genes in the input list), while the color bar indicates -log_10_ (adjusted *P*-value) of the enrichment analysis. (D) Correlation plot comparing log_2_FC values from pathway-level differential expression analysis in both musical stimuli [9] and in TBI *vs.* healthy controls. (E) Upset plot showing the intersection among gene sets identified across the different datasets included in the analysis.

In the same line, ORA of these two clusters of genes (**Table S3**) detected dysregulated pathways involved in neurological conditions. Pathways enrichment analysis from the cluster of genes showing opposite expression changes in TBI and musically stimulated donors may represent processes dysregulated in TBI that are compensated by the effect of the musical stimuli (**Figure 2C**, **Table S3**). Notably, these genes were enriched in processes involved in other neurodegenerative diseases, such as Parkinson’s or Huntington’s disease. In addition, demyelination and axonal polyneuropathy emerged as other pathways related to brain-related disorders. Genes with gene expression changes showing the same direction in music and TBI regulate pathways involved in mitochondrial and metabolic dysfunction (**Figure 2C**, **Table S3**).

Correlation between global activation patterns of pathways altered by musical stimuli and those differentially dysregulated in TBI revealed commonalities between results from the gene-level and pathway-level analysis (**Figure 2D; Table S4**). In fact, mitochondrial and neurological-related processes were identified in both groups of analysis.

Finally, we validated the set of 154 genes commonly impacted by music and altered in TBI using an independent cohort of patients. This cohort included serum miRNA expression profiles of chronic and acute TBI patients as well as age-matched healthy controls. First, we inferred target genes of the differentially expressed miRNA between TBI cases and controls in both acute and chronic TBI patients separately. Then, we intersected each list of target genes with the set of 154 genes and identified 38 common genes that were both affected by the musical stimuli and showed either direct or miRNA-mediated dysregulation due to the TBI condition (**Figure 2E**; **Table S5**). Pathways analysis using these 38 genes revealed significant enrichment in the cGMP-PKG signaling pathway involving genes such as *KCNMA1*, *PPP3CA*, *PRKG1,* and *VDAC1*.

### Co-expression modules and hub genes associated with traumatic brain injury

To explore transcriptional co-regulation patterns associated with TBI, we conducted a *WGCNA* using 4,376 genes selected for high variance across samples. The analysis identified 24 distinct co-expression modules, each assigned a unique color and named by a hub gene representing the most connected gene within the module (**Figure S1**; **Table S6**). Module-trait correlation analysis revealed six modules with moderate to strong correlations with TBI (absolute correlation |r| > 0.5, *P*-value < 0.05); **Table S6**. Among these, the *TXNIP* module showed the strongest negative correlation with TBI (r = –0.69, *P*-value = 0.0048), followed by the *JSRP1* module (r = –0.65, *P*-value = 0.0093); **Figure 3A**. In contrast, four modules were positively correlated with TBI: *PIPOX* (r = 0.59, *P*-value = 0.0216), *NEK2* (r = 0.56, *P*-value = 0.0284), *ALG13* (r = 0.57, *P*-value = 0.0267), and *AQP10* (r = 0.53, *P*-value = 0.0407); **Figure 3A**.

**Figure 3.**
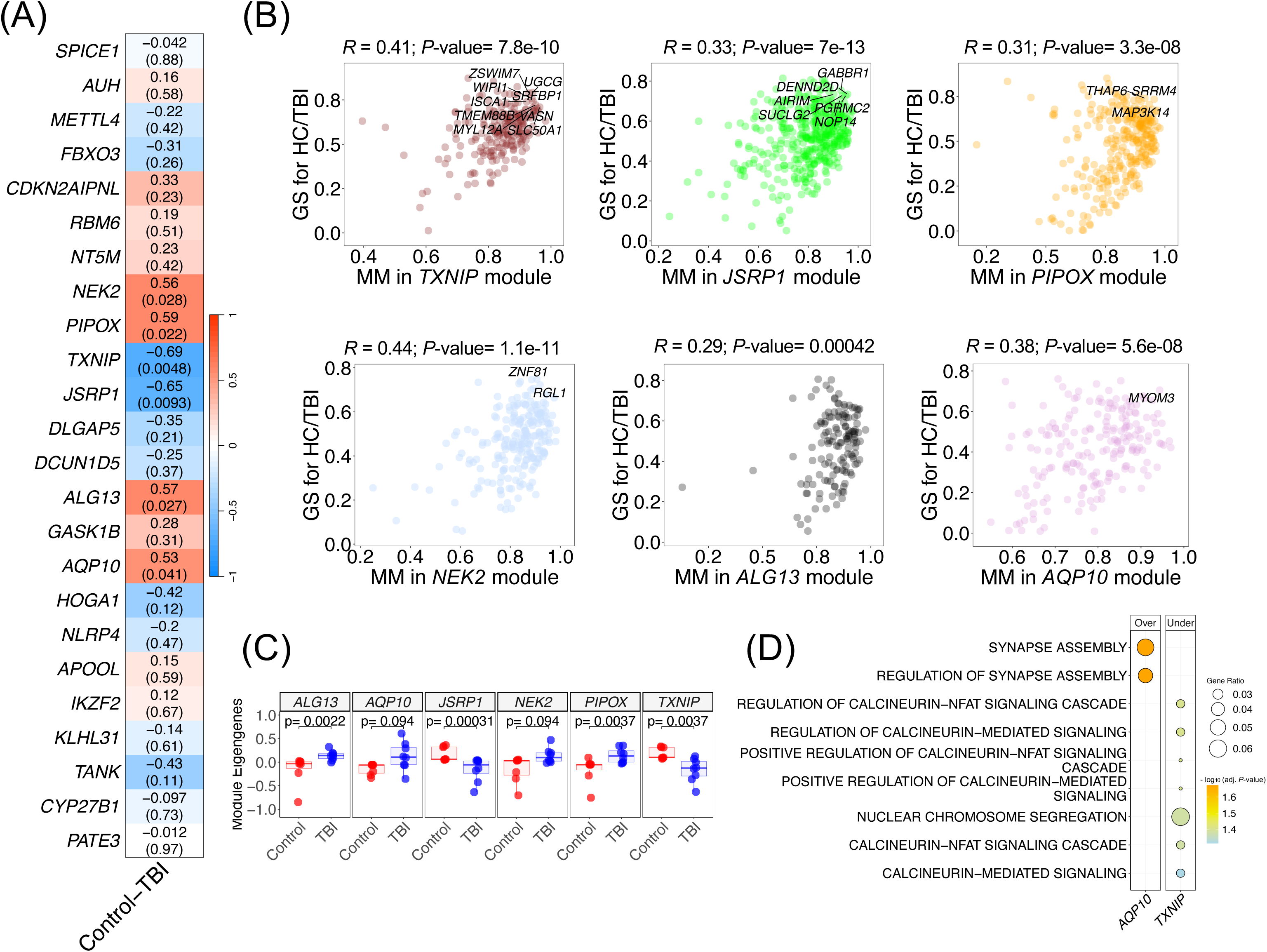
Transcriptomic module associations with TBI status in ASD subjects. (A) Heatmap showing correlation values between modules of co-expressed genes and TBI status (Control *vs*. TBI). Modules are labeled using their respective hub gene names. The upper values indicate the correlation coefficient, while the *P*-values are provided in parentheses. (B) Scatter plot comparing Module Membership (MM) and gene-level correlation with TBI for genes in significantly associated modules. Gene names are shown only for those with MM > 0.9 and correlation > 0.7. (C) Boxplots illustrating differences in individual eigengene expression between Control and TBI samples for statistically significant modules. (D) Enriched biological processes identified from the most significant modules, with modules named according to their hub genes.

Comparison between module membership (MM) and gene significance (GS) revealed a strong and statistically significant correlation across the six modules significantly associated to with TBI (**Figure 3B**), indicating that genes with high intramodular connectivity also tend to exhibit strong individual associations with TBI. The eigengene values of individual samples closely reflected the direction and magnitude of the module-trait correlations, reinforcing the differential co-expression patterns observed between controls and TBI patients (**Figure 3C**). Specifically, modules positively correlated with the condition showed increased eigengene expression in TBI, while negatively correlated modules demonstrated a reduction in eigengene values in TBI. Modules with the lowest *P*-value in the module-trait correlation also showed higher significant differences in eigengene values between controls and TBI patients.

Functional enrichment analysis identified pathways associated with the *TXNIP* and *AQP10* modules. The *TXNIP* module, with a negative correlation with TBI, contained genes with significant over-representation of terms related to calcineurin-NFAT signaling and nuclear chromosome segregation. Notably, the expression of genes from the *TXNIP* module indicates an overall downregulation of this module in TBI, suggesting suppression of calcineurin-dependent pathways (**Table S7; Figure 3D**). Conversely, the *AQP10* module, which displayed a moderate positive correlation with TBI, was enriched for gene ontology terms related to synapse assembly and regulation of synaptic structure (**Table S7; Figure 3D)**.

In order to identify potential music-responsive gene targets within these co-expression modules, we examined whether genes shown to be affected by musical stimuli in healthy controls (derived from [9]) are also driver genes in the significant modules correlated with TBI. We found that between 4-6% of the genes within each module overlapped with the gene set affected by musical stimuli (**Table S8**; **Figure 4**). Notably, a subset of these genes may play a central role in the modules (high MM values), and also showed high correlation values with TBI, demonstrating not only a functional relevance in TBI pathology but also in music-related molecular processes.

**Figure 4.**
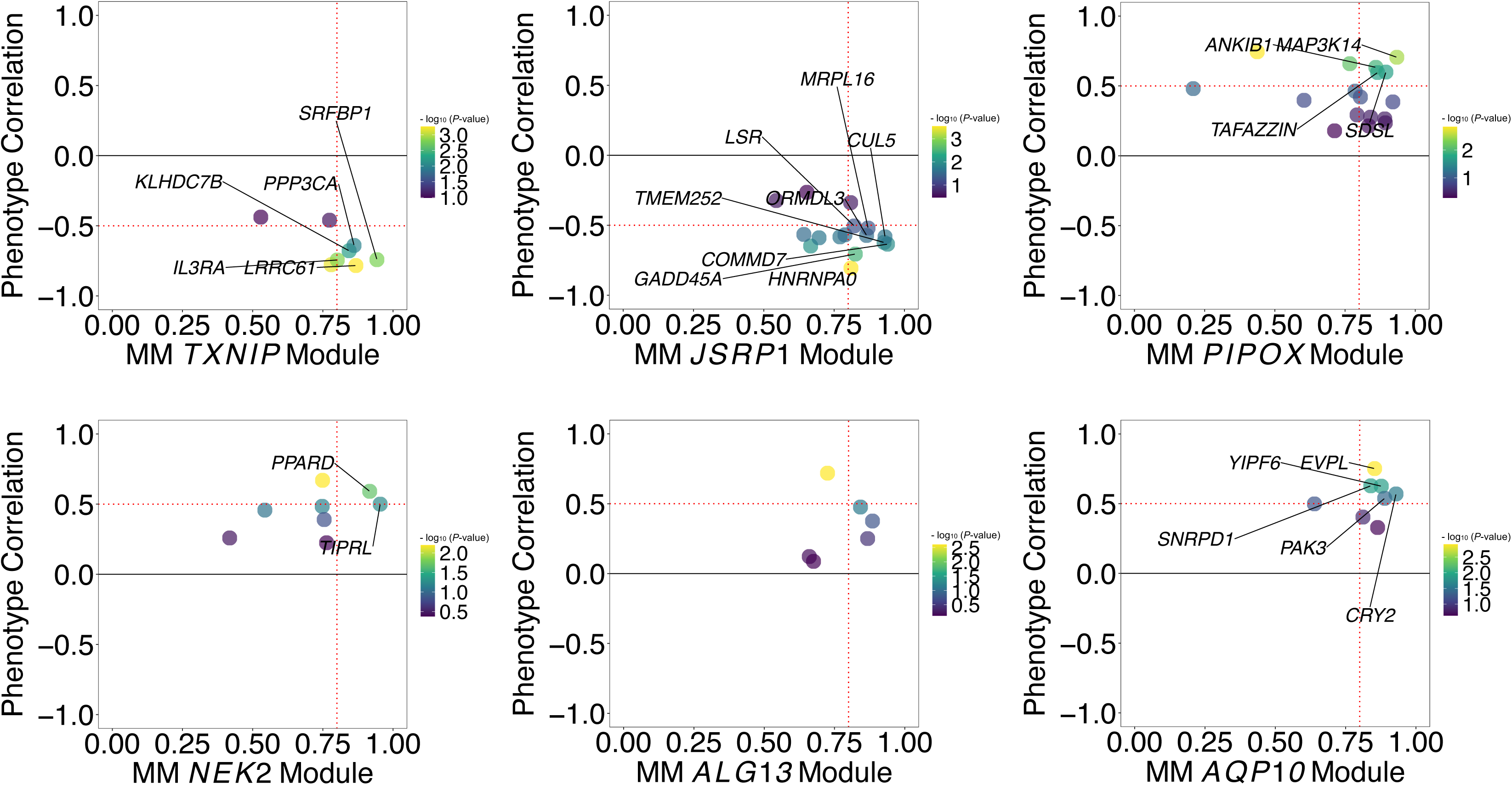
Differentially expressed genes (DEGs) in healthy controls after musical stimulation that are members of TBI-associated modules. Only genes with a Module Membership (MM) > 0.8 (indicated by the vertical dotted line) and an absolute correlation with the phenotype > 0.5 (indicated by the horizontal dotted line) are labelled.

### DNA methylation and traumatic brain injury

Following quality control and normalization of β-values derived from Illumina MethylationEPIC BeadChip array data, the analysis focused specifically on CpG sites annotated to a set of 154 genes previously identified as both responsive to music exposure and altered in TBI through transcriptomic data. Out of the 587,987 CpG sites that passed quality control, 4,764 CpGs mapped to 141 of the 154 genes were retained for statistical testing. A series of independent t-tests comparing individuals with and without a self-reported history of TBI identified 219 CpG sites across 92 genes as significantly differentially methylated (*P*-value < 0.05) (**Table S9**).

To assess the discriminative potential of these 219 CpGs, PCA was performed. The PCA plot (**Figure 5A**) revealed a clear separation between individuals with and without a history of TBI. The first two principal components accounted for 20.8% and 5.8% of the total variance, respectively. Further, hierarchical clustering of methylation levels at the 219 significant CpG sites (**Figure 5B**) showed distinct epigenetic profiles between the two groups, supporting the robustness of the findings. Boxplots of β-value distributions for the 10 most significant CpG sites (**Figure 5C**) highlighted significant differential methylation in genes such as *CCT2, CLEC5A, CRB3, LRRC61, NADK, NUAK1, PPARD, PSMD8, SERBP1,* and *SH2D3C,* with several sites exhibiting either consistent hypermethylation or hypomethylation in the TBI group. Pathways enrichment analysis of the 92 differentially methylated genes reveals significant overrepresentation in five KEGG pathways (**Figure 5D**), including those associated with “Prion disease”, “Huntington disease”, “cGMP-PKG signaling pathways”, all of which are implicated in neurodegenerative processes. Key genes contributing to these pathways included *PSMD8*, *PP3CA*, *VDAC1*, *COX7C*, *PSMA3*, *TUBB2A*, and *UQCRFS1*.

**Figure 5.**
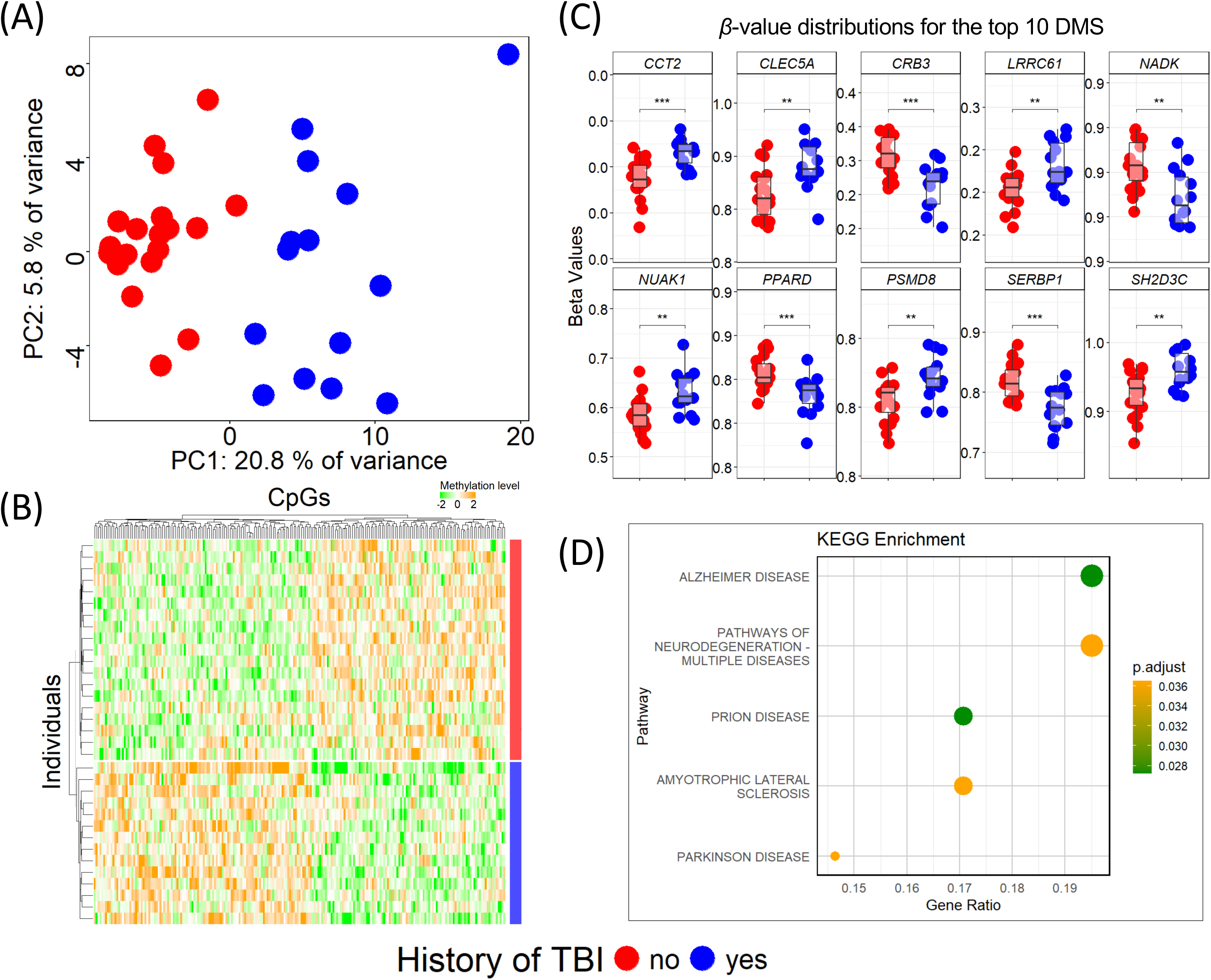
DNA methylation patterns associated with self-reported TBI history in active-duty military personnel. (A) PCA of the 219 CpG sites significantly differentially methylated (*P*-value < 0.05) between individuals with (blue) and without (red) a history of TBI. (B) Heatmap illustrating hierarchical clustering based on the methylation levels at the same 219 CpG sites, showing distinct epigenetic profiles between the two groups. (C) Boxplots displaying β-value distributions for the top 10 differentially methylated CpG sites, annotated to key genes including *CCT2, CLEC5A, CRB3, LRRC61, NADK, NUAK1, PPARD, PSMD8, SERBP1*, and *SH2D3C*. Asterisks indicate levels of statistical significance (*** *P*-value < 0.001, ** *P*-value <0.01). (D) Bubble plot displaying significantly enriched KEGG pathways (adjusted *P*-value < 0.05) identified through over-representation analysis of differentially methylated genes. Point size reflects the gene ratio, and point colour represents the adjusted *P*-value, indicating the strength of statistical significance.

## Discussion

Our recent systematic review and meta-analysis [11] provide strong evidence for the clinical efficacy of music-based interventions in individuals with BD. Consistent improvements were observed across motor, cognitive, communicative, and psychosocial domains. Motor deficits, especially gait and upper limb function, showed notable responsiveness, likely driven by mechanisms such as rhythmic entrainment and auditory-motor coupling that enhance coordination and motivation. Communication gains, particularly in individuals with aphasia, were linked to structured melodic and rhythmic techniques like singing and melodic intonation therapy, which may promote neuroplasticity and functional reorganization in language-related brain regions. Cognitive outcomes were especially robust, with improvements in memory, executive function, and attention, often accompanied by neuroplastic changes observed in frontal and temporal areas. Psychosocial benefits, including mood and social interaction, were also reported, though findings on quality of life (QoL) were less consistent, likely due to methodological variability. Notably, despite some individuals developing amusia post-injury, many, especially those with prior musical training, retained musical abilities, suggesting a potential neuroprotective effect of long-term musical engagement.

Despite the clear clinical potential of music therapy, research into the molecular mechanisms underpinning these therapeutic effects remains scarce. It has been previously reported that music can shape gene expression in a specific manner, highlighting the potential of music-based interventions as an external, non-invasive modulatory factor of the human transcriptome [8-10, 33, 34]. In the context of age-related cognitive disorders, the sensogenomics project highlighted the compensatory role of music in modulating genes and biological processes disrupted in these patients, suggesting a beneficial effect of music for these neurodegenerative conditions [8-10]. However, despite the growing interest in understanding the molecular mechanisms underlying TBI, studies investigating transcriptomic and methylation profiles in patients with TBI remain limited. Furthermore, research examining the effects of music on gene expression and epigenomic regulation in this population remains virtually nonexistent. To address this gap, we adopted a proof-of-concept approach by integrating transcriptomic and epigenomic data from patients with severe TBI with gene expression profiles previously reported in healthy individuals following musical stimulation, as described by Gómez-Carballa et al. [10].

Our analysis identified a subset of 154 DEGs that were commonly affected by both TBI and musical exposure. Among these were key regulators such as *STK35*, *HNRNPA0*, *MT-CYB*, and *MT-ND2*, all of which are implicated in mitochondrial function, synaptic plasticity, and neuroinflammatory responses, processes central to TBI pathophysiology. Interestingly, genes that showed opposite expression patterns between TBI-related profiles and those induced by music were enriched in pathways associated with neurodegenerative disorders, including Parkinson’s and Huntington’s diseases, as well as demyelination and axonal polyneuropathy. These processes also intersect with known mechanisms of TBI and BD, suggesting that music might counteract maladaptive gene expression patterns related to these conditions [35-38]. Both demyelination and axonal damage are commonly observed in TBI but are also hallmarks of many neurodegenerative diseases [39]. Demyelination, characterized by the loss of the myelin sheath and oligodendrocyte cell death, is a major component of white matter injury and contributes significantly to long-term sensorimotor and cognitive deficits [40]. Conversely, genes exhibiting similar directional changes in both contexts were significantly involved in mitochondrial and metabolic processes. These findings reinforce previous results pointing to sustained alterations in synaptic mitochondrial bioenergetics in TBI [41-43]. TBI is characterized by increased neuronal energy demand, ATP depletion, and calcium dysregulation. As a consequence of this mitochondrial dysfunction, purine bases are released into circulation, serving as indicators of systemic mitochondrial impairment [44]. This convergence highlights the pivotal role of mitochondrial impairment across these disorders and suggests that music may modulate key biological pathways disrupted in TBI. It has been shown that TBI also induces metabolic dysfunctions that exacerbate secondary brain damage and are associated with increased risk of mortality in TBI [45]. Hyperammonemia disrupts glutamate homeostasis and mitochondrial activity, contributing to neurotoxicity [46]. Concurrently, lactic acidosis, a marker of anaerobic metabolism, is also a signal of mitochondrial failure and cerebral hypoxia. Both conditions reflect a systemic metabolic crisis that may actively contribute to TBI progression through oxidative stress and mitochondrial impairment [47, 48]. Furthermore, a focused analysis of 38 genes validated in an independent cohort highlighted the cGMP-PKG signaling pathway, particularly involving *KCNMA1*, *PPP3CA*, *PRKG1*, and *VDAC1*, as a potential mediator of music’s neurobiological effects. cGMP-PKG pathway, which is active in neurons, has a well known role in synaptic plasticity and memory consolidation and may represent a molecular target through which music-based interventions exert therapeutic benefits [49-51]. Overall, these pieces of evidence from gene expression studies suggest that musical stimuli may influence genes and pathways involved in TBI (and other neurological conditions), modulating gene expression in a compensatory manner.

DNA methylation analysis identified 92 of the 154 common DEGs as differentially methylated between individuals with and without a history of TBI. Although the most affected genes did not overlap with the top 10 DEGs identified in the transcriptomic analysis, many are known to play critical roles in neurological function and disease. Notably, *CCT2* [52] and *CLEC5A* [53] have been implicated in neurodegenerative disorders such as Alzheimer’s disease (AD). *NUAK1* is linked to neurodevelopmental and psychiatric conditions, including ASD and schizophrenia [54]. *PPARD* is involved in mood regulation and neurogenesis [55], while *SERBP1* has been associated with ischemic stroke risk through specific genetic variants [56]. Pathway analysis further supported findings from the transcriptomic studies, highlighting involvement of the cGMP–PKG signaling pathway as well as other pathways related to neurodegenerative disorders. Together, these findings provide preliminary but compelling evidence that genes modulated by musical stimulation also exhibit differential DNA methylation patterns in individuals with a history of TBI. This suggests a potential epigenetic intersection between music-related gene regulation and TBI-associated molecular changes. By focusing on a shared set of genes from prior transcriptomic analyses, this study highlights specific CpG sites and gene targets that may play a meaningful role in the pathology and recovery of TBI.

Co-expression analysis of TBI and control samples revealed six distinct modules significantly associated with TBI, suggesting a functional relevance of these modules in this condition. Interestingly, some hub genes of the significant modules have been previously linked to TBI pathology. The hub gene of the most significantly correlated module, with Thioredoxin Interacting Protein (*TXNIP*) gene as hub gene, is involved in reactive oxygen species (ROS)-dependent inflammatory response. ROS can upregulate *TXNIP*, which in turn interacts with *NLRP3* to promote activation of the NLRP3 inflammasome, leading to the release of pro-inflammatory mediators [57]. Increased *TXNIP* expression and *NLRP3* inflammasome activation have been reported in the cortex of TBI mice, indicating that *TXNIP* may be a key factor mediating the crosstalk between oxidative stress and neuroinflammation [58, 59]. Activation of *NLRP3* inflammasome via *TXNIP* has also been reported in neurodegenerative conditions such as AD [60, 61]. *De novo* variants in the hub gene of the *ALG13* module, which is a X-linked gene encoding a protein that heterodimerizes with *ALG14*, cause the X-linked congenital disorders of glycosylation type I (ALG13-CDG) [62], a rare human genetic condition that manifests with several different symptoms, including intellectual disability, infantile spasms, and epileptic encephalopathy [63, 64]. Epilepsy is a commonly observed phenotype in ALG13-CDG and presents as an early infantile epileptic encephalopathy [65, 66]. Studies in animal models indicated that *ALG13* may contribute to epilepsy by modulating GABA_A_A receptor function, offering potential insights for therapeutic intervention [67]. Notably, TBI is one of the main causes of acquired epilepsy [68, 69], suggesting that *ALG13* module may represent a functional link between the two conditions. Finally, the hub *AQP10* gene encodes for an aquaporin (aquaporin 10 protein) and, although is mainly an intestinal aquaporin, its function within the CNS has not been investigated yet. Aquaporins are membrane channels that facilitate the transport of water and small solutes. In the CNS, they regulate fluid homeostasis, cerebrospinal fluid dynamics, and metabolic balance, while contributing to neuroprotection under stress. Dysregulation of aquaporins has been implicated in both traumatic and non-traumatic brain injuries, where it may worsen cerebral edema, metabolic dysfunction, and neuroinflammation [70, 71].

Functional enrichment analysis provided additional insights into the potential roles of these modules in TBI, pointing to altered synaptic remodeling mechanisms as well as dysregulated neuroimmune signaling and inflammatory regulation. The calcineurin–NFAT signalling pathway demonstrated a dual role in TBI. Its dysregulation contributes to neuroinflammation, excitotoxicity, and cell death, yet it also plays roles in recovery and repair. Studies in animal models of TBI reported a key role for astrocytic calcineurin/NFAT in disrupting synaptic recovery post-TBI [72]. However, findings on *NFAT4* regulation remain controversial. While one study observed reduced *NFAT4* protein levels in rats following TBI [73], several others have reported robust upregulation of *NFAT4* in activated astrocytes in response to acute TBI [72, 74].

Our indirect identification of module genes potentially regulated by music revealed that some of the most centrally connected genes within each TBI module are likely to play a regulatory or mechanistic role in mediating the transcriptomic response to musical exposure. This convergence supports the hypothesis that music-responsive hub genes within TBI-associated modules could represent promising therapeutic targets for music-based interventions aimed at modulating neural recovery or plasticity following TBI.

While these results offer promising insights into how music may modulate dysregulated pathways in TBI, they represent an initial and indirect step toward understanding the therapeutic potential of music-based interventions. This analysis provides proof-of-concept, demonstrating the feasibility and value of integrating transcriptomic and epigenomic data to investigate music’s molecular effects. As demonstrated in prior work on AD [9], such integrative, omics-based strategies can provide valuable mechanistic insight into non-pharmacological therapies and support the development of evidence-based, personalized music interventions in neurorehabilitation. However, there is a clear need for future studies that directly assess the impact of music in well-characterized in BD patient cohorts, using standardized music intervention protocols and paired molecular profiling before and after treatment. As shown in AD research [10], such approaches could uncover mechanistic links between music and neural recovery, paving the way for evidence-based applications of music therapy in neuropsychiatric care.

Our comparative approach has enabled the identification of shared and divergent gene expression patterns between musical stimulation and TBI, highlighting potential candidates through which music might exert neurobiological effects in this condition. These preliminary findings offer compelling proof-of-concept evidence that musical exposure can modulate key molecular pathways implicated in TBI, potentially counteracting maladaptive gene expression changes through mechanisms involving mitochondrial function, synaptic signaling, and epigenetic regulation. Supporting this, our co-expression analysis identified TBI-associated gene modules enriched for inflammatory and synaptic processes, some of which included hub genes also responsive to music, suggesting possible mechanistic points of convergence. Our strategy provides a valuable first step towards elucidating the potential of music-based therapies in modulating gene expression, which could contribute to neural recovery mechanisms in TBI. However, while promising, this remains an indirect approach, and future research must focus on prospective studies in clinically characterized BD cohorts, incorporating standardized music-based interventions and paired omics profiling before and after treatment.

## Acknowledgements

The authors would like to express their appreciation to the study investigators of the Sensogenomics network (sensogenomics.com; Sensogenomics Working Group [see Annex]), as well as the nursery and laboratory service at the Hospital Clínico Universitario de Santiago de Compostela, for their invaluable dedication and support. This research project was made possible through the access granted by the Galician Supercomputing Center (CESGA) to its supercomputing infrastructure. The supercomputer FinisTerrae III and its permanent data storage system have been funded by the Spanish Ministry of Science and Innovation, the Galician Government, and the European Regional Development Fund (ERDF). This work was supported by: *i*) GAIN IN607B 2020/08 and IN607A 2023/02, and EUTERPE_adn (Programa de Cooperación Interreg-VI POCTEP; Ref. 0313_EUTERPE_ADN_1_E) (to A.S.), IIN607A2021/05 (to F.M.-T.) and IN677D 2024/06 (to A.G.-C.), and *ii*) Consorcio Centro de Investigación Biomédica en Red de Enfermedades Respiratorias (CB21/06/00103; to A.S. and F.M.-T.). AG-C is supported by the Miguel Servet contract (CP23/00080), funded by the Instituto de Salud Carlos III (ISCIII) and co-funded by the European Union. The funders were not involved in the study design, collection, analysis, interpretation of data, the writing of this article, or the decision to submit it for publication.

## Supplementary Material

**Table S1**. DEGs between TBI and healthy controls obtained from the study GSE254880 (adjusted *P*-value <0.05).

**Table S2**. DEGs after musical stimuli from the study by Gómez-Carballa et al. [9] that are also differentially expressed in TBI patients when compared to healthy controls. *P*-value from TBI *vs*. healthy controls was calculated using a Wilcoxon test. **Table S3**. Over-representation analysis of two clusters of genes reported in **Table S1**. GO: Gene ontology; KEGG: Kyoto Encyclopedia of Genes and Genomes.

**Table S4**. Differentially regulated pathways after musical stimuli from the study by Gómez-Carballa et al. [9] that are also differentially regulated in TBI patients when compared to healthy controls.

**Table S5**. Genes altered by the musical stimuli were also dysregulated in the TBI condition (left). Significantly dysregulated pathway obtained from over-representation analysis using the set of 38 genes altered by the musical stimuli, also dysregulated in the TBI condition (right).

**Table S6**. Correlation between co-expression gene modules and TBI status (Controls *vs.* TBI). Corresponding *P*-values for each correlation are also provided.

**Table S7**. Clustered over-representation analysis of significantly correlated modules. GO: Gene Ontology terms.

**Table S8**. Genes modulated by musical stimulation in healthy controls (as reported in Gómez-Carballa et al. [9]) that are also part of the modules significantly correlated with TBI.

**Table S9**. List of 219 significant CpG sites (*P*-value < 0.05) annotated to 92 of the 154 genes previously identified as both responsive to music exposure and altered in traumatic brain injury (TBI) based on transcriptomic data. Delta beta indicates the absolute difference in methylation levels between individuals with and without a history of TBI.

Figure S1. Dendrogram of genes and co-expression modules derived from the *WGCNA* analysis, represented by different colors.

### Sensogenomics Working Group

Antonio Salas Ellacuriaga – PI; Federico Martinón-Torres – PI; Laura Navarro Ramón – Coordinator *GenPoB/GenVip -Instituto de Investigación Sanitaria (IDIS) (alphabetic order)*

Alba Camino Mera, Albert Padín Villar, Alberto Gómez Carballa, Alejandro Pérez López, Alicia Carballal Fernández, Ana Cotovad Bellas, Ana Isabel Dacosta Urbieta, Narmeen Mallah, Ana María Pastoriza Mourelle, Ana María Senín Ferreiro, Andrés Muy Pérez, Antía Rivas Oural, Antonio Justicia Grande, Antonio Piñeiro García, Anxela Cristina Delgado García, Belén Mosquera Pérez, Blanca Díaz Esteban, Carlos Durán Suárez, Carmen Curros Novo, Carmen Gómez Vieites, Carmen Rodríguez-Tenreiro Sánchez, Celia Varela Pájaro, Claudia Navarro Gonzalo, Cristina Serén Trasorras, Cristina Talavero González, Einés Monteagudo Vilavedra, Estefanía Rey Campos, Esther Montero Campos, Fernando Álvez González, Fernando Caamaño Viñas, Francisco García Iglesias, Gloria Viz Rodríguez, Hugo Alberto Tovar Velasco, Irene Álvarez Rodríguez, Irene García Zuazola, Irene Rivero Calle, Iria Afonso Carrasco, Isabel Ferreirós Vidal, Isabel Lista García, Isabel Rego Lijo, Iván Prieto Gómez, Iván Quintana Cepedal, Jacobo Pardo Seco, Jesús Eirís Puñal, José Gómez Rial, José Manuel Fernández García, José María Martinón Martínez, Julia Cela Mosquera, Julia García Currás, Julián Montoto Louzao, Lara Martínez Martínez, Laura Navarro Marrón, Lidia Piñeiro Rodríguez, Lorenzo Redondo Collazo, Lúa Castelo Martínez, Lucía Company Arciniegas, Luis Crego Rodríguez, Luisa García Vicente, Manuel Vázquez Donsión, María Dolores Martínez García, María Elena Gamborino Caramés, María Elena Sobrino Fernández, María José Currás Tuala, María Martínez Leis, María Soledad Vilas Iglesias, María Sol Rodriguez Calvo, María Teresa Autran García, Marina Casas Pérez, Marta Aldonza Torres, Marta Bouzón Alejandro, Marta Lendoiro Fuentes, Miriam Ben García, Miriam Cebey López, Montserrat López Franco, Nour El Zahraa Mallah, Narmeen Mallah, Natalia García Sánchez, Natalia Vieito Perez, Patricia Regueiro Casuso, Ricardo Suárez Camacho, Rita García Fernández, Rita Varela Estévez, Rosaura Picáns Leis, Ruth Barral Arca, Sandra Carnota Antonio, Sandra Viz Lasheras, Sara Pischedda, Sara Rey Vázquez, Sonia Marcos Alonso, Sonia Serén Fernández, Susana Rey García, Vanesa Álvarez Iglesias, Victoria Redondo Cervantes, Vanesa Álvarez Iglesias, Wiktor Dominik Nowak, Xabier Bello Paderne, Xabier Mazaira López

### Nursing volunteers (alphabetic order)

Alejandra Fernández Méndez, Ana Isabel Abadín Campaña, Ana María León Caamaño, Ana María Buide Illobre, Ángeles Mera Cores, Carmen Nieves Vastro, Carolina Suarez Crego, Concepción Rey Iglesias, Cristina Candal Regueira, Dolores Barreiro Puente, Elvira Rodríguez Rodríguez, Eugenia González Budiño, Eva Rey Álvarez, Fernando Rodríguez

Gerpe, Gemma Albela Silva, Isabel Castro Pérez, Isabel Domínguez Ríos, José Ángel Fernández de la Iglesia, José Cruces Vázquez, José Luis Cambeiro Quintela, José Ramón Magariños Iglesias, Julia Rey Brandariz, Julio Abel Fernández López, Luisa García Vicente, Manuel González Lito, Manuel González Lijó, Manuela Pérez Rivas, Margarita Turnes Paredes, María Aurora Méndez López, María Begoña Tomé Arufe, María Campos Torres, María del Carmen Baloira Nogueira, María del Carmen García juan, María Esther Moricosa García, María Luz Chao Jarel, María Martínez Leis, María Mercedes Jiménez Santos, María Salomé Buide Illobre, María Victoria López Pereira, Mercedes Jorge González, Mercedes Isolina Rodríguez Rodríguez, Miren Payo Puente, Natalia Carter Domínguez, Olga María Reyes González, Pilar Mera Rodríguez, Purificación Sebio Brandariz, Salomé Quintáns lago, Yolanda Rodríguez Taboada, María Pereira Grau.

### Other volunteers (alphabetic order)

Alba Arias Gómez, Alejandro Moreno Díaz, Ana Arca Marán, Astro González Guirado, Brais García Iglesias, Carlos Sánchez Rubín, Carmen Otero de Andrés, Clara Pérez Errazquin Barrera, Claudia Rey Posse, Cristina Rojas García, Eduardo Xavier Giménez Bargiela, Elena Gloria Morales García, Fabio Izquierdo García Escribano, Gabriel Guisande García, Jaime López Martín, Lara Pais Ramiro, Lucía Rico Montero, Luís Estévez Martínez, Manuel Estévez Casal, María Aránzazu Palomino Caño, María Rubio Valdés, Marisol Nogales Benítez, Miryam Tilve Pérez, Nuria Villar Muiños, Pablo Del Cerro Rodríguez, Pablo Pozuelo Martínez Cardeñoso, Salma Ouahabi El Ouahabi, Santiago Vázquez Calvache

