## Supplementary figures and images for "Can music modulate gene expression involved in traumatic brain injury? An integrative transcriptomic and epigenomic proof of concept"

### Figure S1

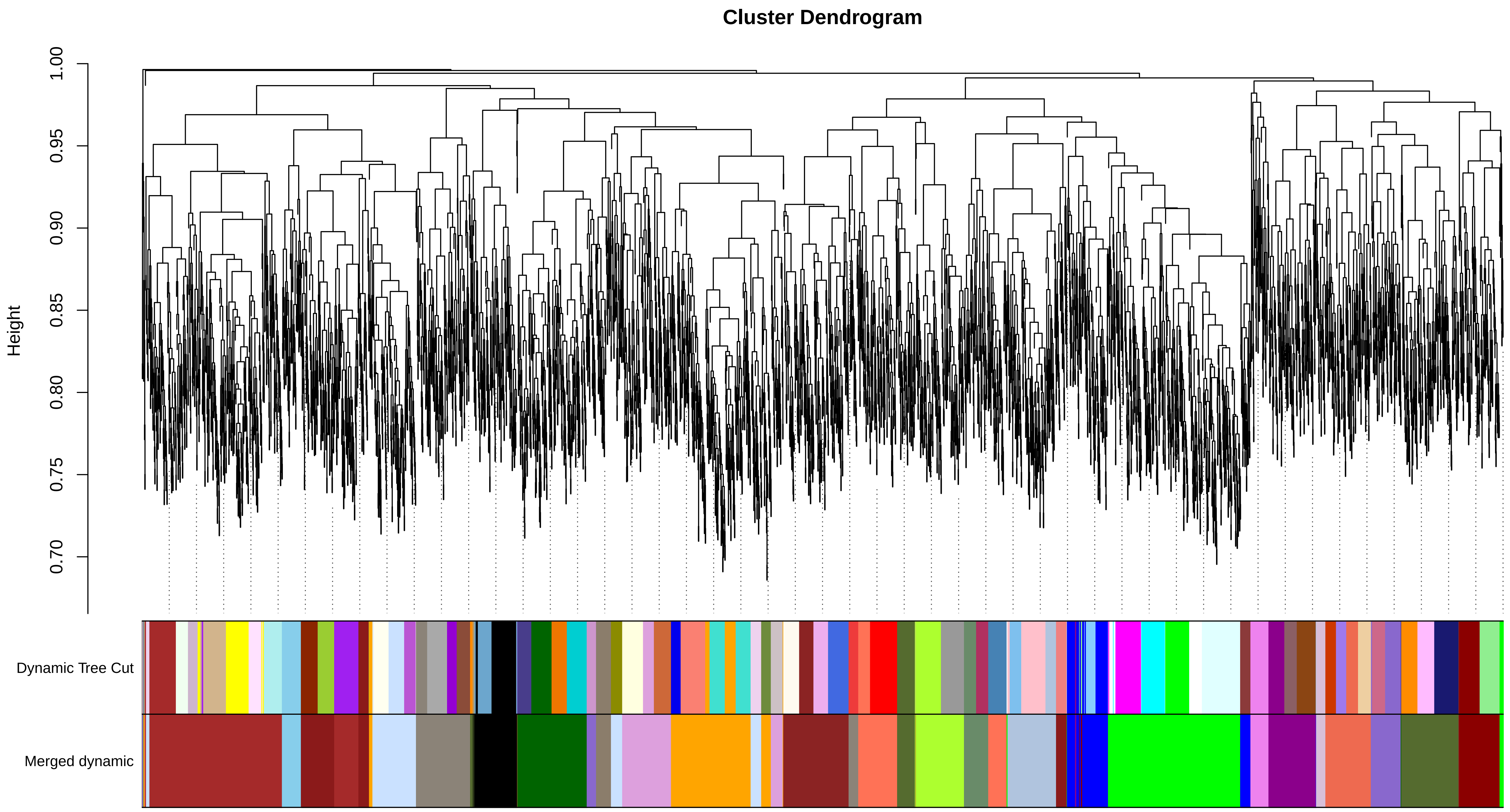
